# Physico-chemical characterization of single bacteria and spores using optical tweezers

**DOI:** 10.1101/2022.10.24.513442

**Authors:** Daniel P.G. Nilsson, Unni Lise Jonsmoen, Dmitry Malyshev, Rasmus Öberg, Krister Wiklund, Magnus Andersson

**Affiliations:** Department of Physics, Umeå University, 901 87 Sweden; Department of Paraclinical Sciences, Faculty of Veterinary Medicine, Norwegian University of Life Sciences, 1433 Norway; Umeå Center for Microbial Research (UCMR), 901 87 Sweden

**Keywords:** endospores, Raman spectroscopy, metabolic activity, adhesion, pili, CaDPA

## Abstract

Spore-forming pathogenic bacteria are adapted for adhering to surfaces, and their endospores can tolerate strong chemicals making decontamination difficult. Understanding the physico-chemical properties of bacteria and spores is therefore essential in developing antiadhesive surfaces and disinfection techniques. However, measuring physico-chemical properties in bulk does not show the heterogeneity between cells. Characterizing bacteria on a single-cell level can provide mechanistic clues usually hidden in bulk measurements. This paper shows how optical tweezers can be applied to characterize single bacteria and spores, and how physico-chemical properties related to adhesion, fluid dynamics, biochemistry, and metabolic activity can be assessed.

## 1. INTRODUCTION

A pathogenic microorganism’s ability to aggregate, colonize a surface, or infect a host relies on its physico-chemical properties.^1^ These properties are also essential for bacterial fitness and survival in harsh environments. Some pathogenic bacterial species within the genera *Bacillus* and *Clostridium* are experts in surviving tough environments thanks to their ability to sporulate (turn from vegetative cell to endospore) in response to unfavorable conditions. They form bacterial endospores (spores) that are highly resistant to environmental challenges compared to their vegetative cell form.^2^ Spores can survive high heat (including boiling water), radiation, and chemical exposure to common antimicrobial agents,^3^ and their resistance increases when adhered to a surface.^4^ Pathogenic spore agents are therefore problematic in many areas of society, from contaminating equipment in food production facilities^5^ and hospitals,^6^ to spores of *Bacillus anthracis* being highly potent agents for biological warfare.^7^

Part of what makes pathogenic spores problematic are their surface properties. Bacteria and spores have a sophisticated nano-machinery, being able to express proteinaceous fiber-like surface organelles that help facilitate adhesion to host surfaces. These structures, commonly denoted pili or fimbriae in bacteria, are widespread within both Gram-negative and Gram-positive bacteria and spores.^8–10^ In Fig. 1A we show a SEM micrograph of a *Bacillus cereus* spore expressing pili (green arrows). Pili are comprised of numerous subunits (pilins) that are formed via different pathways and subsequently compose pili of varying architecture and functionality. In vegetative bacteria, pili are involved in adhesion, biofilm formation, cell aggregation, host cell invasion, motility, and secretion and uptake of DNA and proteins.^11^ In addition, the presence of pili affects the bacterium’s surface characteristics. The intrinsic physical properties of adhesion pili are crucial for infection since bacteria expressing mutant pili with comprised physical properties have been shown to lose their capacity to cause infection.^12^ Understanding the physical properties of pili can help reveal structural features explaining not only pili’s fundamental role and function in attachment but also provide critical targets for new therapeutics to combat and prevent microbes from adhering. However, since pili are nanometer-wide and micrometer-long fibers, sophisticated nanotools are required to quantify their physical properties governed by interactions on the nanometer scale.

**Figure 1.**
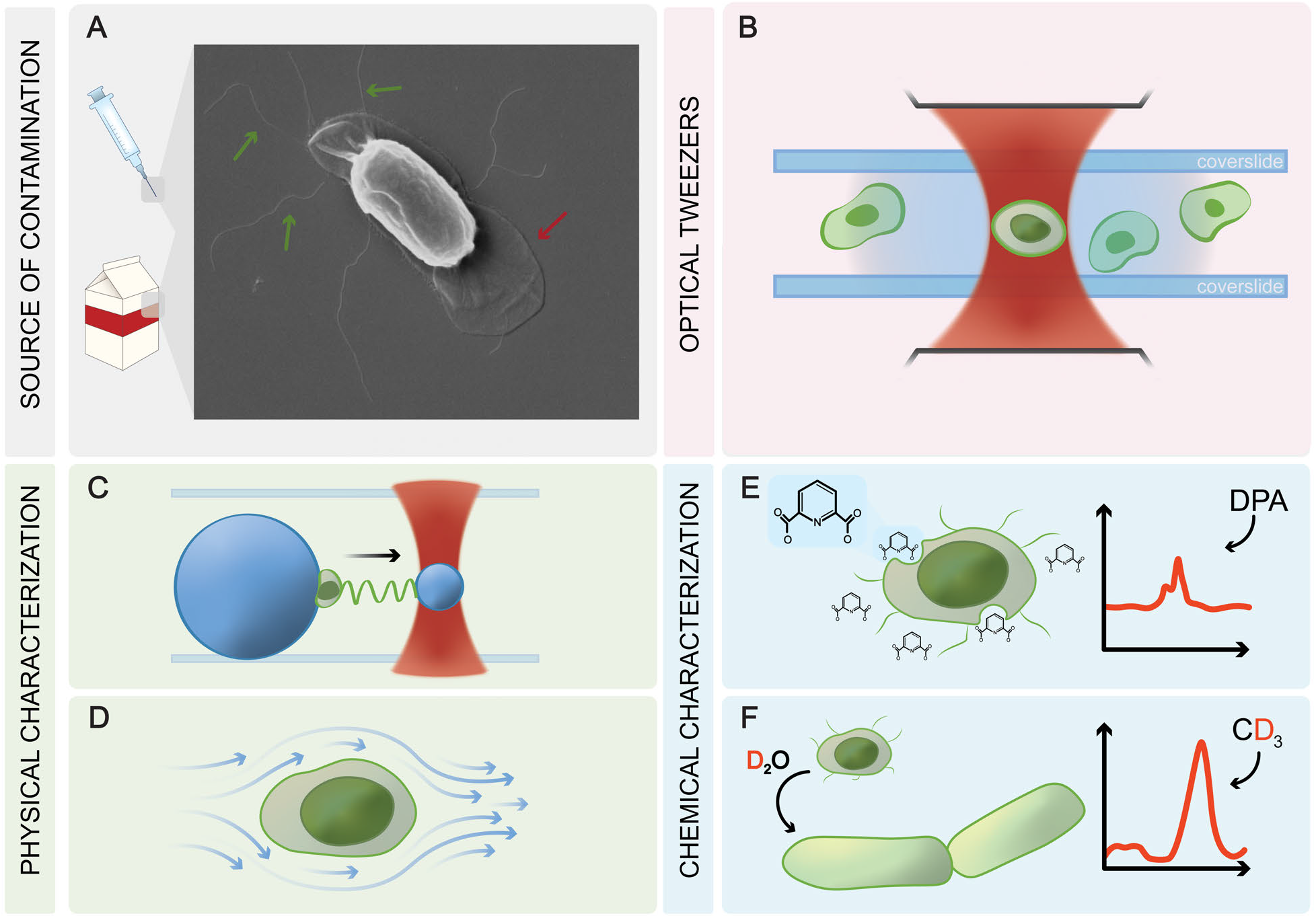
Microorganisms can stick to surfaces and create biofilms, resulting in problems in healthcare facilities and food industries. Spore adhesion is strongly related to physico-chemical properties (pili, exosporium, hydrophobicity, charge, surface structure, etc.). Panel (A) shows a micrograph of a spore with pili (green arrows) and the exosporium (red arrow) clearly visible. (B) Optical tweezers can trap single spores inside a suspension by focusing a laser beam with a microscope objective. (C) By trapping a probe bead, it is possible to measure the mechanical properties of nanofibers, for example pili, by attaching the bead to a pilus and monitoring the bead’s position in the trap. (D) By rapidly moving (oscillating) a trapped spore inside the suspension, its hydrodynamic properties can be measured, and with an applied electric field its electric charge can be measured. The trapping laser also produces inelastically scattered light (Raman scattering) from the trapped spore, which contains information on the biochemical, content making it possible to identify a specific molecule (E). Monitoring the Raman scattered light, also allows non-invasively measuring metabolic activity (F) in the spore by adding a small amount of heavy water.

Bacterial and spore adhesion, as well as resistance to chemical disinfection treatments, also depend on their chemical properties. Therefore, several bulk studies have aimed to characterize properties such as hydrophobicity and zeta potential^4, 13^ in relation to surface adhesion and biofilm formation. However, measuring these properties using population-based methods does not disclose heterogeneities within a population. Cell divergence and heterogeneity between cells within the same strain and environmental condition have received more attention in the last decade.^14–16^ This is especially critical for spore-forming bacteria where large variations in stages of the life cycle and sporulation might occur in the same culture, or biofilm.^15, 17^ Thus, characterizing bacteria on a single-cell level offers insight into cell properties beyond that of bulk measurements and helps provide a better understanding of their pathogenic mechanisms.

There are many ways to study a single cell, but one of the most versatile nanotools are optical tweezers.^18, 19^ Optical tweezers are a technique that makes use of the fact that micrometer sized objects (inorganic as well as biological objects) can be trapped by well focused laser light. It is thereby possible to non-intrusively control and manipulate living biological objects in a fully controlled manner with great precision solely by light. Therefore, optical tweezers can be used to trap, move, and manipulate microspheres, bacteria, and spores, as illustrated in Fig. 1(B). This makes optical tweezers especially suited for measuring receptor-ligand bonds or the mechanical properties of proteins (C), hydrodynamic drag and electric charge of a spore (D), chemical markers (E), and metabolic activity (F) in cells. With optical tweezers it is possible to study the mechanics of cell membranes, surface organelles, track the chemical content of a single cell over time, and non-invasively detect and characterize pathogens.^19, 20^

Optical tweezers work by focusing laser light to a diffraction-limited (smallest possible) spot using a microscope objective. If a cell is close, it interacts with the light and is pulled into the trap by gradient optical forces.^18^ When pulling on a trapped particle, the force that is required increases with the displacement in the trap, similar to a regular spring. Thus, knowing the trap stiffness (spring constant) allows us to measure external forces acting on a particle by measuring its displacement. Further, a trapped particle can be moved in all three dimensions by either moving the focal spot or by translating the sample chamber (keeping the trap stationary). The sample chamber is made from two microscope glass slides, separated by a thin liquid layer. Moving a trapped particle back-and-forth in the liquid, makes it possible to measure hydrodynamic drag forces.

In addition to light’s capability to trap a particle, light will also interact with the molecular bonds in the particle. This results in inelastically scattered light that have either gained or lost energy during the interaction. This inelastically scattered light is called Raman scattering.^21^ By using a spectrometer to analyze the Raman scattered light, it is possible to extract information about the particle’s molecular composition. The resulting Raman spectrum is a unique fingerprint of the particle that can be used to identify the chemical composition of the particle and follow how it changes with time non-invasively.^22^ Thus, there are many biological applications where optical tweezers can be used to measure the physical and chemical properties of single bacteria or spores. In this work, we describe a few of these applications.

## 2. APPLICATIONS OF OPTICAL TWEEZERS

### 2.1 Moving, sorting, and isolating single spores with light

To conduct measurements on a single cell, the cell must first be separated from the larger population. Since optical tweezers can be applied to both trap micro-sized objects and move them through a liquid at tens of micrometers per second, they provide a unique way to separate, sort, and isolate single cells to study heterogeneity.^23, 24^ For example, to assess the laser power threshold for causing spore damage in an optical trap, Malyshev et al. exposed spores to different laser powers. After trapping, the spores were deposited on the coverslide and organized in a pattern to keep track of each individual spore. The coverslide with deposited spores could thereafter be evaluated using high resolution SEM imaging and the visual damage could be correlated with prior laser exposure.^25^ Correlation assays, in which cells are first exposed to physical or chemical damage on the single-cell level and later studied using imaging methods (SEM, fluorescence, AFM, etc.), allows for new types of mechanistic studies.

### 2.2 Measuring physical properties using force spectroscopy

Optical tweezers can also be used to measure forces by adding a detector suitable for the laser light wavelength. When light interacts with a particle in the trap, some light is scattered from the particle and some is transmitted. The interference of the light produces a diffraction pattern that gives a direct measure of the particle’s displacement in the trap. Via calibration, it is possible to derive the absolute force acting on the particle with high resolution. Thanks to both the high spatial and temporal resolution it is possible to study a multitude of physical properties and assess structural, kinetic, and cell parameters, for example, adhesion pili mechanics, molecular motor-driven flagellar rotation, and the hydrodynamic diameter of cells.

#### 2.2.1 Forces on bacteria and spore appendages

There is a broad repertoire of surface pili found on bacteria with distinct structural and functional properties.^11^ Variation of properties can be investigated using optical tweezers that allow quantitative measurements where force is applied and measured in the range from sub-piconewton to hundreds of piconewton. These forces are what is needed to separate receptor-ligand interactions, unfold proteins, extract tethers from cell membranes, and characterize nanopolymers.^20, 26^ In particular, optical tweezers force measurements have proven exceptionally useful in measuring the physical properties of bacterial and spore pili.^27–29^

A simple and cheap assay to measure the physical and mechanical properties of cell-surface expressed nanofibers, is to trap a bacterium (or spore) using optical tweezers and position the bacterium onto a poly-l-lysine coated 10 μm microsphere, that is immobilized to the coverslide.^30^ Thus, the microsphere serves as a mount for the particle under study, see Fig. 2(A). With the bacterium attached to the larger bead, a smaller probe bead is subsequently trapped and moved close to the bacterium to attach a pilus, in general, the longest expressed. Separating the bacterium and probe bead, tensile force is applied, and the force-extension/contraction response is measured with sub-piconewton force and nanometer spatial resolution. Examples of force-extension curves of a P pilus expressed by *Escherichia coli* and a S-Ena pilus of a *B. cereus* spore are shown on Fig. 2(B,C). Two types of appendages have been described on the *B. cereus* NVH 0075/95 spores, denoted S-Ena and L-Ena appendages.^31^ The S-Ena fibers are longer and thicker, while the L-Enas are shorter and thinner. Force extension and retraction curves using the setup described illustrates significant differences in mechanisms and functionality between the S-Ena of *B. cereus* and the *E. coli* P pili. The force data indicate that the P pilus can be unwound, and with a constant extension force they extended significantly (6x its initial length). During extension the quaternary structure of the fiber undergoes several conformational transitions. The S-Ena, on the other hand, does not unwound at these forces, and shows a limited axial extensibility and no conformational changes to its structure. These two examples highlight the variety of mechanistic differences that force measurements are able to reveal. By further applying physical models to the data, for example the worm-like-chain, intrinsic physical parameters can be assessed such as persistence length, contour length, stretch modulus, spring constant and flexural rigidity.

**Figure 2.**
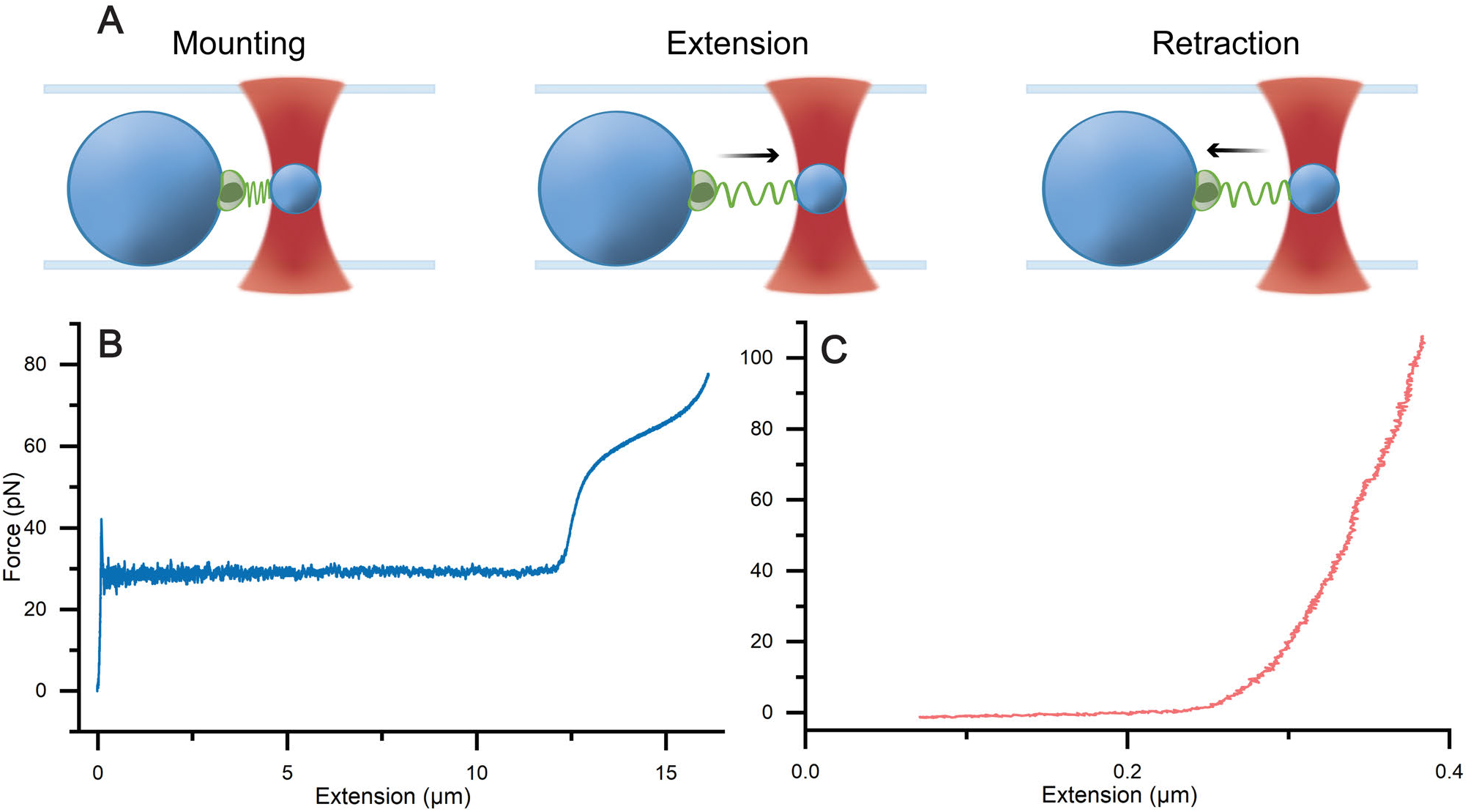
Force-extension experiments on bacteria using optical tweezers. Panel (A) shows the process of an optical tweezers force measurement. (Left) A pili expressing bacterium or spore is trapped and mounted to a microsphere, and subsequently a probe bead is trapped and attached to a pilus. (Middle) The probe bead and microsphere are separated to extend the pilus. (Right) After the pilus has been extended, the beads can be brought back together to relax the pilus. Panel (B) shows the force response of a helical P pilus expressed from an *E. coli* bacterium. The curve shows a constant force with extension. Panel (C) displays the force response of a S-Ena pilus expressed by a *B. cereus* spore. This force response simply increases with distance and never reaches a plateau, making the extension much shorter.

#### 2.2.2 Measuring motility of single bacteria

Bacteria face numerous challenges in their natural environments and have developed different approaches to both sense chemicals and swim away from danger. For example, many bacteria swim using flagellar rotation driven by molecular motors at the base.^32^ Some bacteria, such as *Bacillus subtilis* or *E. coli*, even express multiple flagellar filaments that bundle together in the so-called run sequence. Nevertheless, this bundle can be disturbed, resulting in a tumble sequence in which some flagella rotate at different speeds or change their directions.^33^ Understanding how bacteria switch between the run and tumble event, as well as how the molecular motor changes its frequency in the presence of nutrients and chemicals, is relevant in chemotaxis studies. However, observing these events on the nanometer scale of free-swimming bacteria is challenging using bulk methods. Therefore, optical trapping of free-swimming bacteria opens a new door to characterizing such events.

By acquiring the scattered light from the optically trapped bacterium, the rotation motion of the bacterium’s body and the flagellum can be determined. A schematic of this is shown in Fig. 3(A), in which the bacterial rotation Ω and flagellum *ω* rotates in opposite directions. These rotation frequencies are clearly visible in a power spectrum graph, as illustrated in Fig. 3(B). Using optical tweezers assays, several important research questions have been solved related to bacterial motility. For example, how flagellar rotational variation of *B. subtilis* depends on changes in pH or the nutrient concentration in the fluid;^34^ how dynamic properties such as power and the thrust of *E. coli* relates to the angular velocity of the motor;^35^ and the importance of elastic properties of flagella in *E. coli* and *Streptococcus* bacteria to work as propellers, as well as to form bundles.^36^

**Figure 3.**
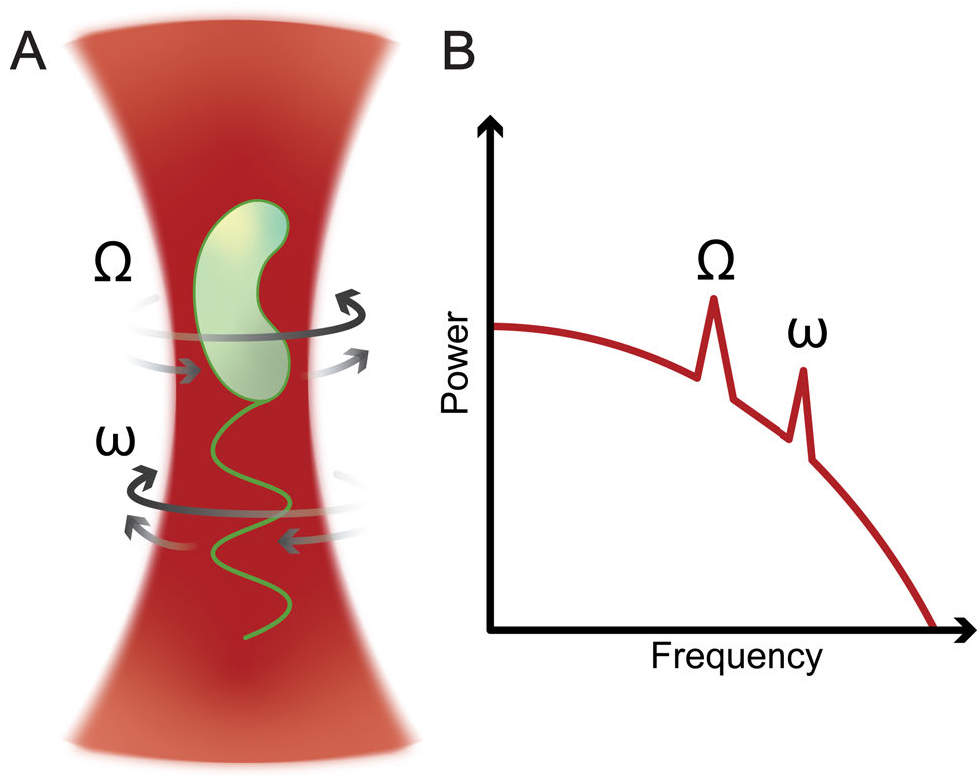
Panel (A) illustrates an optically trapped bacterium, in which the body and flagellum rotates in opposite direction. Panel (B) shows how the corresponding power spectrum can reveals the rotation frequencies. By changing the fluid environments (pH, nutrients, chemicals, etc.) this method allows for direct assessment of the nanomachinery that rotates the flagellum.

#### 2.2.3 Hydrodynamic drag of spores

Endospores are, in general, hydrophobic and strongly adhere to a variety of surfaces, including stainless steel, glass and plastics. This makes spores a major contamination hazard in food processing environments, particularly strains of *B. cereus*. These bacteria are also problematic since they produce biofilms that facilitate attachment and protect vegetative cells and spores from fluid forces and disinfection chemicals.^37, 38^ Since both vegetative bacteria and spores often are exposed to fluid forces in their environments, and there is large heterogeneity in their surface properties, assessing surface properties on the single cell level can reveal mechanistic details that help us better understand their ability to attach to surfaces and each other.

Optical tweezers can measure surface properties by trapping a single cell and oscillating the sample chamber so that the cell is exposed to a fluid drag force from the liquid, see Fig. 4A. From this, it is possible to obtain the cell’s hydrodynamic drag coefficient and calculate its effective diameter. The optical tweezers method also benefits from using very small sample volumes (microliters). It can also, by its sensitivity, observe the effect of extracellular physiological variations, for example, the expression of surface organelles. For instance, oscillating optical tweezers have previously been used to quantify the degree of surface pili expressed on *E. coli* bacterial cells.^39^ The study showed that highly piliated cells experience more than a 2-fold effective diameter (higher drag force) compared to bald cells. The implication of expressing “too” many pili was studied using a motility assay and supports that a large effective diameter can be negative in fluid environments since the swimming capability is significantly impaired and bacteria with many pili tend to form more aggregates.^40^

**Figure 4.**
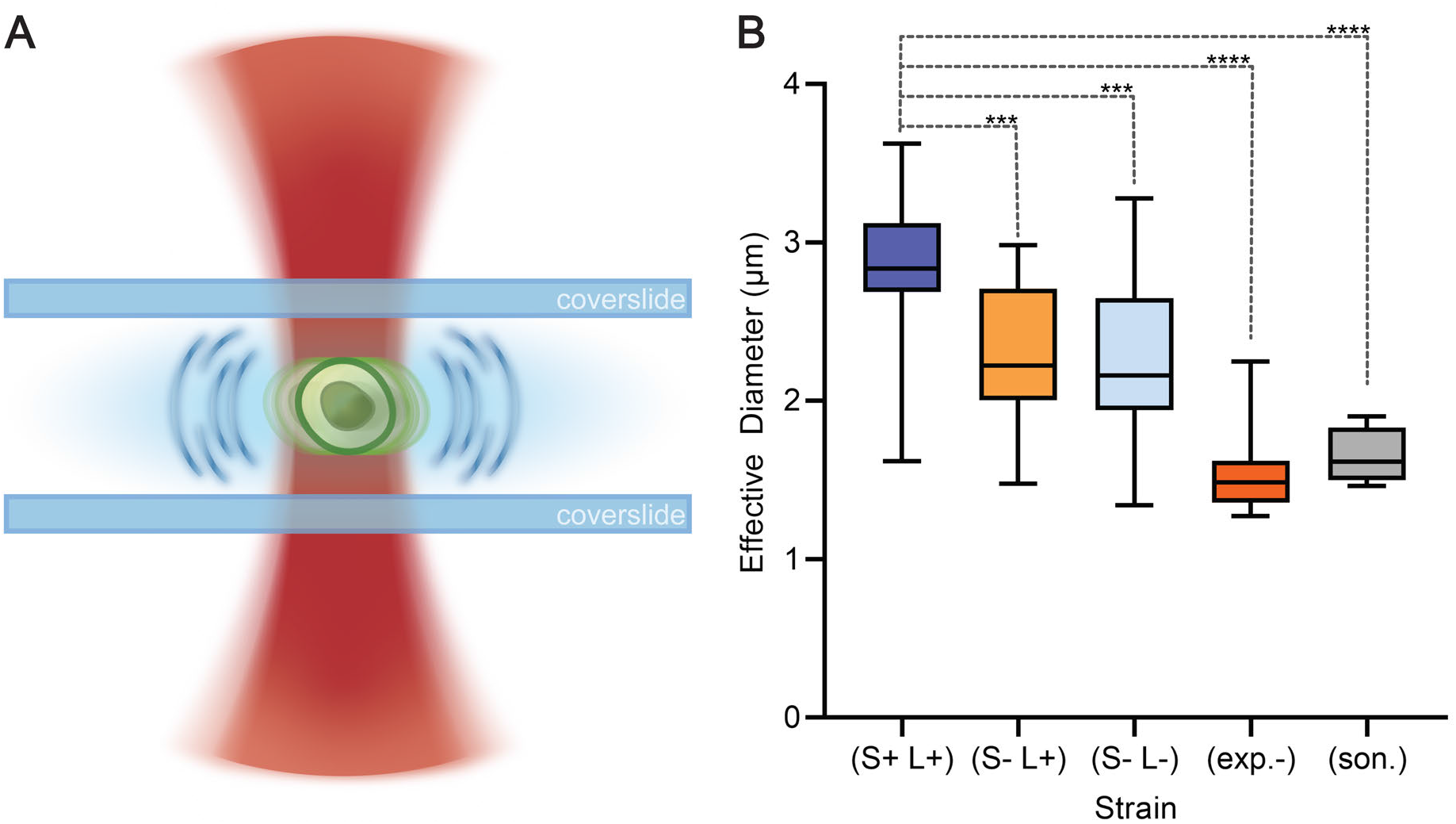
Measuring hydrodynamic drag of a spore with optical tweezers. Panel (A) illustrates how the trap can be oscillated rapidly to expose the spore to a fluid drag force. By measuring the displacement of the spore in the trap the effective hydrodynamic diameter can be calculated. Panel (B) shows effective hydrodynamic diameter of wild type spore (S+ L+), spores with only L-Ena pili (S-L+), spores lacking S-Ena and L-Ena pili (S-L-), spores lacking an exosporium (exp.-) and sonicated spores (son.). Boxes cover the 25-75 percentiles in the data distribution (with the median as a solid line) and the whiskers cover the 5-95 percentile. Stars indicate the statistical significance of the difference compared to the wild type spores, either p ≤ 0.001 (***) or p ≤ 0.0001 (****).

Surface appendages can also affect the drag experienced by spores. A similar procedure has been used to investigate the effect of pili and the exosporium on hydrodynamic drag on *B. cereus* NVH 0075/95 endospores, which expresses both S-Ena and L-Ena appendages. An example of the significant role these surface organelles have on the drag force is shown in Fig. 4B. There, wild type spores (S+ L+) are compared with knock out mutant spores depleted of S-Enas (S-L+), both types of Enas (S-L-), and spores depleted of their exosporium (exp.-).^29^ Additionally, a (S-L-)-strain with its expsoporium mechanically removed by sonication is also included (son.) The results illustrate the sensitivity of using optical tweezers to quantify the difference in drag force (effective diameter) between wild type spores and mutants. The oscillating optical tweezers method can also be taken one step further to simultaneously measure the hydrodynamic drag coefficient and the electric charge, as shown by Pesce et al., when they trapped *B. subtilis* spores. By employing this method, they were able to distinguish the electric charge between wild type spores and spores of isogenic mutant strains that lacked or had a defective outer coat layer.^41^

### 2.3 Characterization of chemical properties using Raman spectroscopy

Optical tweezers can also be used to characterize chemical properties of single bacteria and spores. Such measurements provide information on the status and viability of bacteria in response to different treatments. Having a cell trapped, changes in chemical composition inside the cell can be followed using Raman spectroscopy. When light interacts with a molecule in the trapped particle, some light scatters inelastically off the molecules, gaining or losing energy depending on the energy of the molecular vibrations. This process is called Raman scattering and can be used to acquire Raman spectra, which give information on the molecular bonds present. Raman scattering has the advantage of relatively narrow bands compared to fluorescence and absorbance measurements, making the technique very specific in distinguishing between molecules. Another major advantage is that water is not Raman active, so aqueous samples can be measured without noise from the solvent. Raman scattering signals are weak, so intense light is needed for these measurements and, therefore, optical trapping, where laser light is highly focused on an particle, is very well suited for Raman spectroscopy. The resulting technique, laser tweezer Raman spectroscopy (LTRS) allows for single particle measurements, reducing the noise both from surrounding media and from the surface of the sample chamber since the trapped particle can be moved into the center of the sample channel during a measurement. LTRS has been used in several applications characterizing and monitoring chemical properties and changes within bacterial spores.^42–44^

#### 2.3.1 DPA release from single spores

Bacterial spores are dormant until environmental conditions become more suitable. Then, they can turn back into vegetative cells via a process called germination. Tracking the germination process is important in studies on spore biochemistry and spore resistance to chemicals. During germination, or if damaged by toxic chemicals or high mechanical stress, bacterial spores will release dipicolinic acid (DPA). DPA is abundantly found in calcium chelate form (CaDPA) in the spore core, making up as much 25 % of the spore dry weight. This makes it an excellent biomarker for spores when using Raman spectroscopy.^45, 46^ Thus, by using LTRS, spores and the germination process can be tracked using the Raman peak for CaDPA at 1017 cm^-1^.^47^ In addition, it is possible to follow chemical changes in single bacterial spores as they are exposed to various decontamination process,^48^ as summarized in Fig. 5A. As an example, Fig. 5B shows a time-series of the release of CaDPA from a single spore. Note that this release, when initiated is a rapid process, taking place within the span of a minute, see spectra between 45-46 minutes. Further, it was shown with this technique that the DPA release from chemically exposed *Bacillus thuringiensis* spores is faster when exposed to near-infrared light.^44, 49^ This shows that although spores are highly resilient, photo-chemical effects can arise when spores are exposed to both chemicals and light.

**Figure 5.**
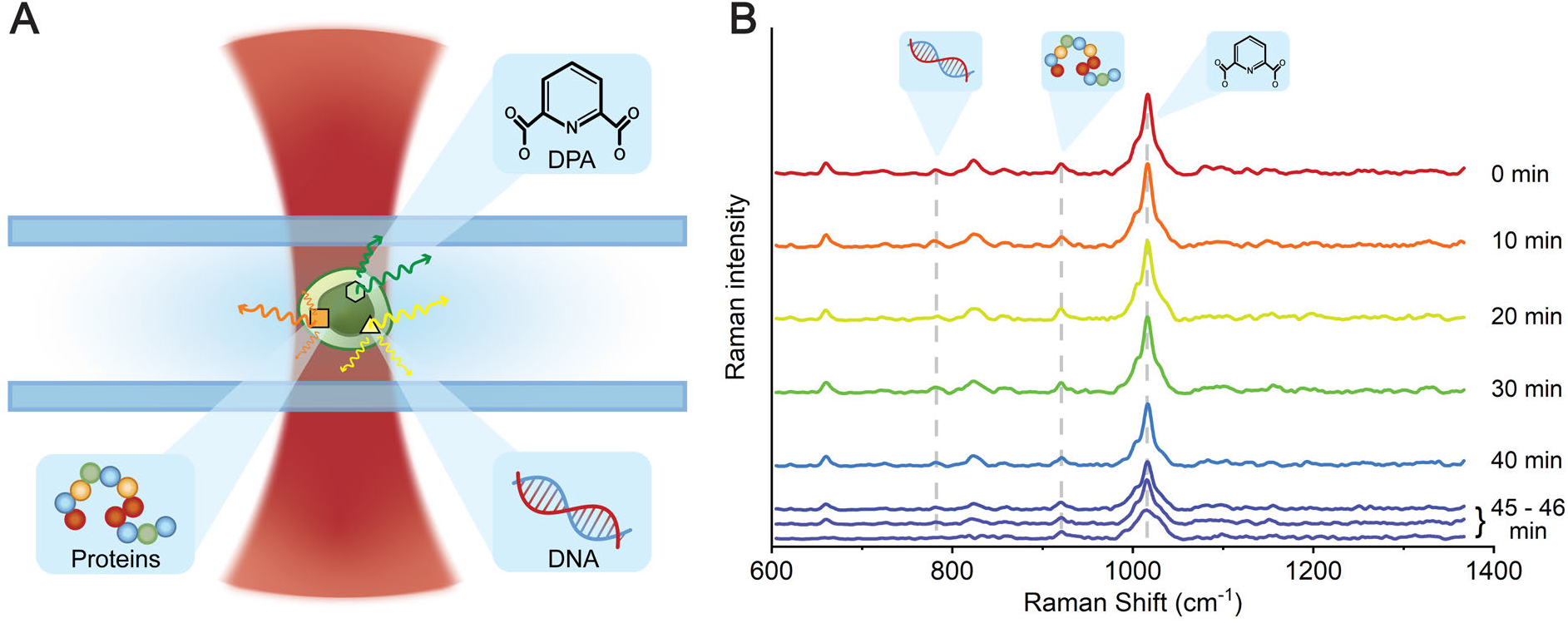
Measuring chemical properties of a single cell using laser tweezers Raman spectroscopy (LTRS). Panel (A) shows that we can use Raman spectroscopy to measure chemical components such as CaDPA, DNA, proteins and lipids when trapping a single spore or bacterium. Panel (B) shows that the loss of CaDPA from a single spore can be tracked as a rapid decrease in intensity of the 1017 cm^-1^ peak (at 45-46 min for this particular spore).

#### 2.3.2 Impact of chemicals on spores

Understanding the effect of chemicals on spores is essential in developing robust decontamination and disinfection protocols. Similarly to tracking DPA release, LTRS can track chemical changes in spores during chemical-based decontamination processes. Earlier studies have used Raman spectroscopy to look at the release of DPA from spores^50^ and the mechanism of common disinfection agents.^51^ Although, these are done as bulk studies and information on how individual spores respond is lost. LTRS offers, again, the advantage of measuring and selecting individual spores,^52^ and have been used on several occasions to investigate spore inactivation in response to disinfectants.^21, 53^ For example, Zhang et al. used LTRS to measure the effectiveness of wet-heat inactivation of *Bacillus* spores.^54^ Further, LTRS has also been used to track chemical changes in *B. thuringiensis* spores as they were exposed to chlorine dioxide, sodium hypochlorite, and peracetic acid.^44^ By following Raman peaks at 782 cm^-1^ and 1017 cm^-1^ during the disinfection process, the amount of DNA and CaDPA, and thus the release of these chemicals from the spore over time, could be quantified. Using this method, it is possible to evaluate the spore damage caused by these chemical treatments and thus evaluate their efficiency.

#### 2.3.3 Metabolic activity of single cells

The possibility to track the metabolic activity of cells in real-time is advantageous as it presents a rapid way to monitor a bacterium’s response to antimicrobials, chemicals, and environmental changes. A simple and low-cost method to track metabolic activity of single bacteria and spores during outgrowth, is to use LTRS in combination with a small amount (about 30%) of heavy water (deuterated water). Water is used as a reactant in cell metabolism, such as the TCA cycle and is gradually incorporated into cell proteins, lipids and DNA. When water is substituted for heavy water, the incorporation of deuterium results in C-D bonds. These bonds can be detected using Raman spectroscopy as a broad band centered at ~2190 cm^-1^. This band is in a quiet Raman region, so the magnitude of this band is a direct indicator of the cumulative metabolic activity in the cell to this point. This method has been used to track metabolic activity and viability of bacterial cells,^55–57^ including cells that appear inactivated, but remain metabolically active.^58^ We have also shown this method to work on monitoring the germination and outgrowth of spores,^59^ as summarized in Fig. 6.

**Figure 6.**
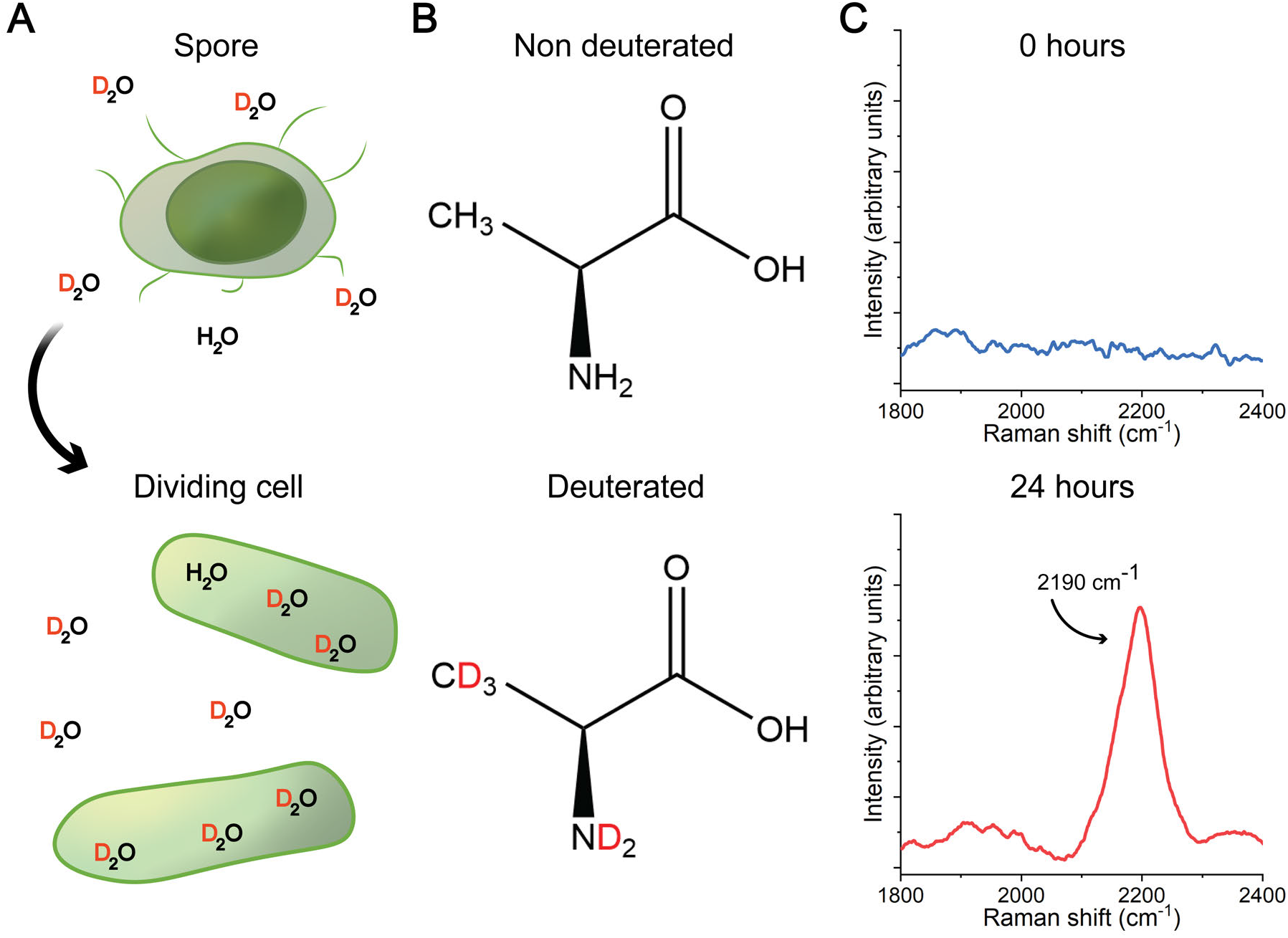
Metabolic activity of germinating *B. cereus* (A) can be tracked using a heavy water assay. Water is taken up and incorporated into proteins and lipids during metabolic activity (B), and the resulting C-D bonds from heavy water can be seen as a new broad Raman band centered at 2190 cm^-1^ (C).

Metabolic analysis using LTRS have several advantages. First, with minimal sample preparation needed the method is rapid and cheap. Second, heavy water its non-toxic (up to ~50 % can be added in buffer solution) to the cells studied. Third, it is versatile and applicable to a wide range of bacterial and spore samples and can be used on all samples that are metabolically capable, including samples directly from the field. Finally, when investigating bacterial viability its quick with no culturing needed.

## 3. CONSIDERATIONS

### 3.1 Instrumental limitations and possible solutions

Even though optical tweezers are powerful nanotools, it is important to consider limitations when designing experiments. To generate a strong trap, an optical tweezers setup needs a high-numerical aperture objective (preferably >1.2) and a stable laser source (intensity and pointing stability). Between the objective and the sample, an immersion medium is used to reduce reflections. Oil is commonly used; however, this limits the trapping distance inside the sample making manipulation and force measurements deep into the sample difficult. A water immersion objective fixes this issue, but water evaporates much faster which limits the experimental time, making long-term measurements more difficult. However, by using a micro dispenser offered by microscope manufactures this issue can be resolved. When preparing samples, it is also important to find a suitable balance between enough particles for the purposes of the experiment and enough to avoid trapping multiple particles.

A setup for measuring forces on the piconewton scale is very sensitive to environmental disturbances. Vibrations and electrical interference can add noise to the data, unless the setup is well isolated. For this, everything from signal filters to acoustic dampening should be considered. When using LTRS, the glass slides that make up the sample chamber can produce a significant background signal, masking Raman peaks of interest. It can be mitigated by using quartz slides, or by moving the trapped particle away from the glass surface. In addition, Raman spectra will periodically acquire cosmic rays, usually appearing as very narrow and intense peaks. This can be circumvented by using multiple accumulations for each spectral acquisition and removing the outlying peaks.

### 3.2 Laser phototoxicity affect cell viability

Optical trapping may be considered non-invasive for inanimate objects,^60^ but it is not the case for living organisms. Even without direct thermal or photo-ablative effects, a 1 W optical trap generates a several MW / mm^2^ spot. It has been shown that even optical traps with a power as low as 3 mW^61, 62^ and doses as low as 0.54 J,^63^ affect cell viability. Trapping introduces DNA damage as well as production of reactive oxygen species (ROS) such as of singlet oxygen,^64, 65^ which in turn affects the function and structural integrity of the cells.^61, 62^ In addition, the strongly focused laser beam (high intensities) in an optical trap can result in that two or more photons are absorbed simultaneously by an atom (non-linear effects). This implies that damage, similar to those from ultraviolet irradiation, can be seen despite trapping the spore with near-infrared light.

These effects are not limited to vegetative bacterial cells. Bacterial spores, despite their resilience, can also be inactivated by optical trapping, see Fig. 7C.^44^ Spores have several mechanisms to protect against ROS and radiation damage, including superoxide transmutases, small acid-soluble proteins and DNA repair mechanisms,^66, 67^ but a sufficiently high dose will deplete them. For a 1064 nm laser, the spores can tolerate approx. 10 J total illumination, with higher energies leading to the spores being unable to germinate, as shown in Fig. 7(A-B). However, laser photo-toxicity effects can be mitigated by using either lower trap power or trapping time. Lower laser power will also reduce the non-linear effects. Finally, the ROS generation can be mitigated by degassing the suspension or adding ROS scavengers like glucose oxidase or protocatechuate dioxygenase to the suspension.^68^

**Figure 7.**
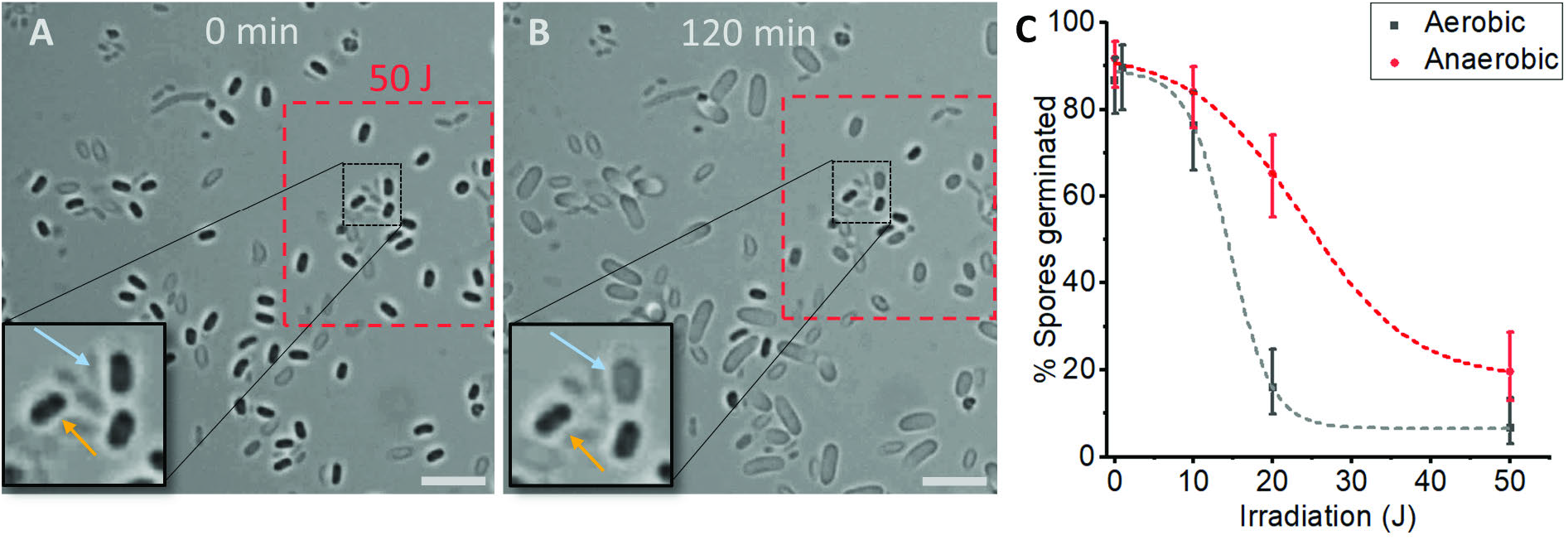
Laser light from optical tweezers can affect the viability of spores. Individual spores in a chosen area (A) were sequentially trapped with at 1064 nm laser (red rectangle). After 120 minutes in nutrient broth, most of the irradiated spores fail to germinate, even as surrounding non-irradiate spores in the same field of view germinated and proceed to grow into the vegetative cells. (B). Some irradiated spores release their DPA (A-B inset, blue arrows), while other retain it (inset, orange arrows), however, both groups fail to grow into vegetative cells. Overall, the spore’s ability to germinate follows a dose-response relationship is and is affected by the oxygenation of water (C).

### 3.3 Raman peak is dependent on solution pH

Raman peaks of complex organic molecules is affected by solution composition. For example, the bond vibrations giving the spectral peaks of these molecules can be affected by the ionic state of the molecule and by chelates bound to it. When comparing spectra in the literature, it is therefore important to evaluate any differences in how solutions were prepared. Also, this implies that reported spectra in the literature can be inconsistent with measurements. As an example, the DPA molecule that typically has a peak at 1000 *cm*^-1^, increases to 1017 *cm*^-1^ when in the chelate (CaDPA) form. Aqueous DPA can have peaks at both 1000 and 1017 cm^-1^ depending on solution pH,^49^ as shown in Fig. 8. Similar studies on biologically relevant compounds such as amino acids reveal similar pH-induced changes in Raman activity,^69, 70^ further showing the importance of knowing and tracking environmental conditions in a measured sample. The change of Raman peaks with pH-condition, is not only something negative, is also provides an opportunity to experimentally derive the pK_a_ of a chemical.^49^

**Figure 8.**
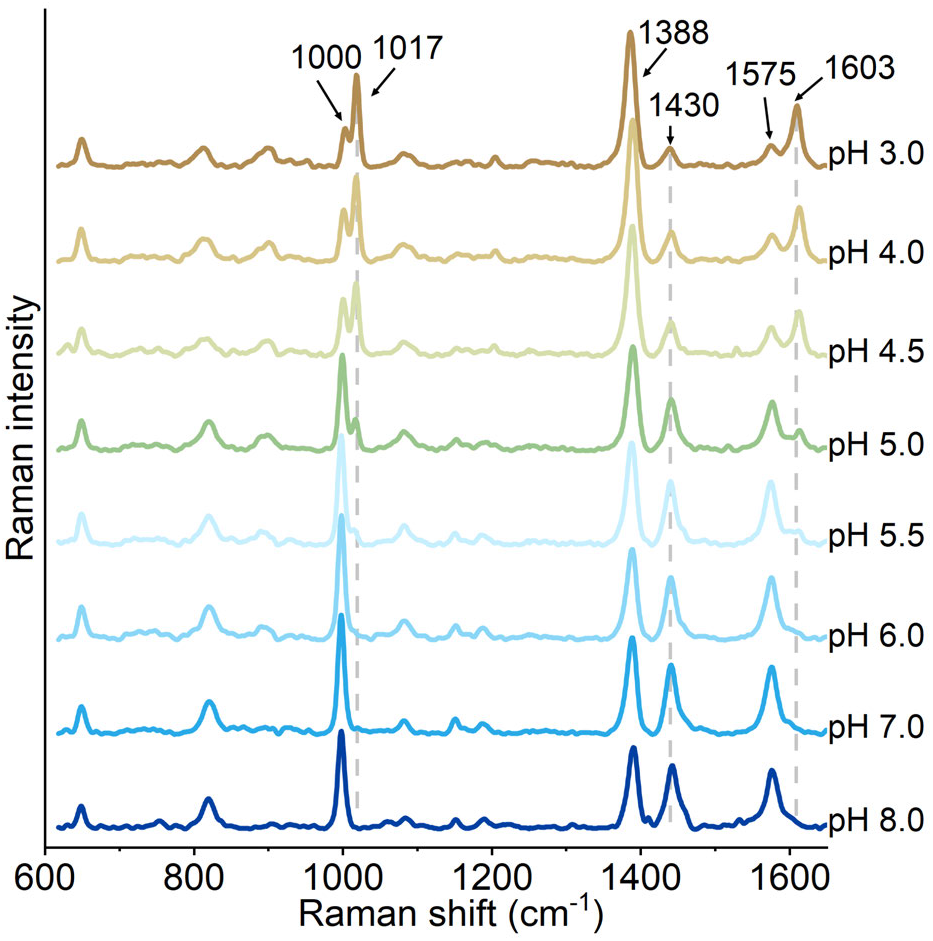
The Raman peaks for pure DPA in aqueous solution shifts as it changes into different ionic forms depending on pH. At pH 3, the major peaks of aqueous DPA are 1017, 1388 and 1603 *cm*^-1^, with smaller peaks at 1000, 1430 and 1575 *cm*^-1^. With increasing pH, these smaller peaks increase in prominence, while the 1017, 1388 and 1603 *cm*^-1^ decrease. At pH ≥ 6, the 1017 and 1603 *cm*^-1^ are no longer detectable.

## 4. CONCLUSION

Since its invention in the 1970s, optical tweezers have proven to be powerful tools in microbiology. The possibility to trap single cells allows for precise micro-manipulation and characterization of physico-chemical properties. These properties include force measurements on single pilus structures and measuring the hydrodynamic drag and electric charge of cells. In addition, the possibility to simultaneously trap and measure Raman signals on particles means that optical tweezers are an effective tool to detect chemical markers in a non-destructive manner. This makes it possible to, among other things, track germination and metabolic activity in real-time.

## ACKNOWLEDGMENTS

This work was supported by the Swedish Research Council (2019-04016); the Umeå University Industrial Doctoral School (IDS); Kempestiftelserna (JCK−916.2); Swedish Department of Defence, Project no. 470-A400821; and Norwegian University of Life Sciences (NMBU).

## Notes

### Competing Interest Statement

The authors have declared no competing interest.

## REFERENCES

[1] Casadevall, A. and Pirofski, L.-a., “Host-Pathogen Interactions: Redefining the Basic Concepts of Virulence and Pathogenicity,” Infection and Immunity 67, 3703–3713 (aug 1999).

[2] Hornstra, L., Leeuw, P., Moezelaar, R., Wolbert, E., de Vries, Y., de Vos, W., and Abee, T., “Germination of Bacillus cereus spores adhered to stainless steel,” International Journal of Food Microbiology 116, 367–371 (may 2007).

[3] Fawley, W. N., Underwood, S., Freeman, J., Baines, S. D., Saxton, K., Stephenson, K., Owens, R. C., and Wilcox, M. H., “Efficacy of Hospital Cleaning Agents and Germicides Against Epidemic Clostridium difficile Strains,” Infection Control & Hospital Epidemiology 28, 920–925 (aug 2007).

[4] Simmonds, P., Mossel, B. L., Intaraphan, T., and Deeth, H. C., “Heat Resistance of Bacillus Spores When Adhered to Stainless Steel and Its Relationship to Spore Hydrophobicity,” Journal of Food Protection 66, 2070–2075 (nov 2003).

[5] Shaheen, R., Svensson, B., Andersson, M. A., Christiansson, A., and Salkinoja-Salonen, M., “Persistence strategies of Bacillus cereus spores isolated from dairy silo tanks,” Food Microbiology 27(3), 347–355 (2010).

[6] Ikram, S., Heikal, A., Finke, S., Hofgaard, A., Rehman, Y., Sabri, A. N., and Økstad, O. A., “Bacillus cereus biofilm formation on central venous catheters of hospitalised cardiac patients,” Biofouling 35, 204–216 (feb 2019).

[7] Manchee, R. J., Broster, M. G., Anderson, I. S., Henstridge, R. M., and Melling, J., “Decontamination of Bacillus anthracis on Gruinard Island?,” Nature 303, 239–240 (may 1983).

[8] Pizarro-Cerda, J. and Cossart, P., “Bacterial Adhesion and Entry into Host Cells,” Cell 124, 715–727 (feb 2006).

[9] DesRosier, J. P. and Lara, J. C., “Isolation and properties of pili from spores of Bacillus cereus,” Journal of Bacteriology 145, 613–619 (jan 1981).

[10] Driks, A., “Surface appendages of bacterial spores,” Molecular Microbiology 63 (feb 2007).

[11] Proft, T. and Baker, E. N., “Pili in Gram-negative and Gram-positive bacteria - structure, assembly and their role in disease.,” Cellular and molecular life sciences: CMLS 66, 613–35 (feb 2009).

[12] Spaulding, C. N., Schreiber, H. L., Zheng, W., Dodson, K. W., Hazen, J. E., Conover, M. S., Wang, F., Svenmarker, P., Luna-Rico, A., Francetic, O., Andersson, M., Hultgren, S., and Egelman, E. H., “Functional role of the type 1 pilus rod structure in mediating host-pathogen interactions,” eLife 7, e31662 (jan 2018).

[13] Jindal, S. and Anand, S., “Comparison of adhesion characteristics of common dairy sporeformers and their spores on unmodified and modified stainless steel contact surfaces,” Journal of Dairy Science 101, 5799–5808 (jul 2018).

[14] Ackermann, M. and Schreiber, F., “A growing focus on bacterial individuality,” Environmental Microbiology 17, 2193–2195 (jul 2015).

[15] Ryall, B., Eydallin, G., and Ferenci, T., “Culture History and Population Heterogeneity as Determinants of Bacterial Adaptation: the Adaptomics of a Single Environmental Transition,” Microbiology and Molecular Biology Reviews 76, 597–625 (sep 2012).

[16] Lin, Y., Briandet, R., and Kovács, Á. T., “Bacillus cereus sensu lato biofilm formation and its ecological importance,” Biofilm 4, 100070 (dec 2022).

[17] Verplaetse, E., Slamti, L., Gohar, M., and Lereclus, D., “Cell Differentiation in a Bacillus thuringiensis Population during Planktonic Growth, Biofilm Formation, and Host Infection,” mBio 6 (jul 2015).

[18] Ashkin, A., Dziedzic, J. M., Bjorkholm, J. E., and Chu, S., “Observation of a single-beam gradient force optical trap for dielectric particles.,” Optics letters 11(5), 288 (1986).

[19] Polimeno, P., Magazzù, A., Iatì, M. A., Patti, F., Saija, R., Esposti Boschi, C. D., Donato, M. G., Gucciardi, P. G., Jones, P. H., Volpe, G., and Maragò, O. M., “Optical tweezers and their applications,” Journal of Quantitative Spectroscopy and Radiative Transfer 218, 131–150 (2018).

[20] Bustamante, C. J., Chemla, Y. R., Liu, S., and Wang, M. D., “Optical tweezers in single-molecule biophysics,” Nature Reviews Methods Primers 1, 25 (dec 2021).

[21] Esposito, A. P., Talley, C. E., Huser, T., Hollars, C. W., Schaldach, C. M., and Lane, S. M., “Analysis of Single Bacterial Spores by Micro-Raman Spectroscopy,” Applied Spectroscopy 57, 868–871 (jul 2003).

[22] Xie, C., Dinno, M. A., and Li, Y.-Q., “Near-infrared Raman spectroscopy of single optically trapped biological cells.,” Optics Letters 27(4), 249–51 (2002).

[23] Keloth, A., Anderson, O., Risbridger, D., and Paterson, L., “Single Cell Isolation Using Optical Tweezers,” Micromachines 9, 434 (aug 2018).

[24] Chapin, S. C., Germain, V., and Dufresne, E. R., “Automated trapping, assembly, and sorting with holographic optical tweezers,” Optics Express 14, 13095 (dec 2006).

[25] Malyshev, D., Robinson, N. F., Oberg, R., Dahlberg, T., and Andersson, M., “Reactive oxygen species generated by infrared laser light in optical tweezers inhibits the germination of bacterial spores,” Journal of Biophotonics 15, 1–7 (aug 2022).

[26] Molloy, J. E. and Padgett, M. J., “Lights, action: Optical tweezers,” Contemporary Physics 43, 241–258 (jul 2002).

[27] Fällman, E., Schedin, S., Jass, J., Andersson, M., Uhlin, B. E., and Axner, O., “Optical tweezers based force measurement system for quantitating binding interactions: System design and application for the study of bacterial adhesion,” Biosensors and Bioelectronics 19, 1429–1437 (jun 2004).

[28] Castelain, M., Duviau, M.-P., Canette, A., Schmitz, P., Loubiere, P., Cocaign-Bousquet, M., Piard, J.-C., and Mercier-Bonin, M., “The Nanomechanical Properties of Lactococcus lactis Pili Are Conditioned by the Polymerized Backbone Pilin.,” PloS one 11(3), e0152053 (2016).

[29] Dahlberg, T., Jonsmoen, U. L., Oberg, R., Malyshev, D., Aspholm, M. E., and Andersson, M., “Endospore pili - flexible, stiff and sticky nanofibers,” (Under Review) Biophysical Journal (2022).

[30] Andersson, M., Fällman, E., Uhlin, B. E., and Axner, O., “Dynamic Force Spectroscopy of E. coli P Pili,” Biophysical Journal 91, 2717–2725 (oct 2006).

[31] Pradhan, B., Liedtke, J., Sleutel, M., Lindbäck, T., Zegeye, E. D., O’Sullivan, K., Llarena, A., Brynildsrud, O., Aspholm, M., and Remaut, H., “Endospore Appendages: a novel pilus superfamily from the endospores of pathogenic Bacilli,” The EMBO Journal 40, 1–16 (sep 2021).

[32] DeRosier, D. J., “The Turn of the Screw: The Bacterial Flagellar Motor,” Cell 93, 17–20 (apr 1998).

[33] Mukherjee, S. and Kearns, D. B., “The Structure and Regulation of Flagella in Bacillus subtilis,” Annual Review of Genetics 48, 319–340 (nov 2014).

[34] Bhat, A. V., Basha, R. A., Chikkaiah, M. D., and Ananthamurthy, S., “Flagellar rotational features of an optically confined bacterium at high frequency and temporal resolution reveal the microorganism’s response to changes in the fluid environment,” European Biophysics Journal 51, 225–239 (apr 2022).

[35] Chattopadhyay, S., Moldovan, R., Yeung, C., and Wu, X. L., “Swimming efficiency of bacterium Escherichia coli,” Proceedings of the National Academy of Sciences 103, 13712–13717 (sep 2006).

[36] Block, S. M., Blair, D. F., and Berg, H. C., “Compliance of bacterial flagella measured with optical tweezers,” Nature 338, 514–518 (apr 1989).

[37] Wijman, J. G., De Leeuw, P. P., Moezelaar, R., Zwietering, M. H., and Abee, T., “Air-liquid interface biofilms of Bacillus cereus: Formation, sporulation, and dispersion,” Applied and Environmental Microbiology 73(5), 1481–1488 (2007).

[38] Kwon, M., Hussain, M. S., and Oh, D. H., “Biofilm formation of Bacillus cereus under food-processing-related conditions,” Food Science and Biotechnology 26(4), 1103–1111 (2017).

[39] Zakrisson, J., Singh, B., Svenmarker, P., Wiklund, K., Zhang, H., Hakobyan, S., Ramstedt, M., and Andersson, M., “Detecting Bacterial Surface Organelles on Single Cells Using Optical Tweezers,” Langmuir 32, 4521–4529 (may 2016).

[40] Cao, L., Suo, Z., Lim, T., Jun, S., Deliorman, M., Riccardi, C., Kellerman, L., Avci, R., and Yang, X., “Role of overexpressed CFA/I fimbriae in bacterial swimming.,” Physical Biology 9(3), 036005 (2012).

[41] Pesce, G., Rusciano, G., Sasso, A., Isticato, R., Sirec, T., and Ricca, E., “Surface charge and hydrodynamic coefficient measurements of Bacillus subtilis spore by optical tweezers.,” Colloids and surfaces. B, Biointerfaces 116, 568–75 (apr 2014).

[42] Chen, D., Huang, S. S., and Li, Y. Q., “Real-time detection of kinetic germination and heterogeneity of single *Bacillus* spores by laser tweezers Raman spectroscopy,” Analytical Chemistry 78(19), 6936–6941 (2006).

[43] Zhang, P., Setlow, P., and Li, Y., “Characterization of single heat-activated Bacillus spores using laser tweezers Raman spectroscopy,” Optics Express 17, 16480 (sep 2009).

[44] Malyshev, D., Dahlberg, T., Wiklund, K., Andersson, P. O., Henriksson, S., and Andersson, M., “Mode of Action of Disinfection Chemicals on the Bacterial Spore Structure and Their Raman Spectra,” Analytical Chemistry 93, 3146–3153 (feb 2021).

[45] Kočiŝová, E. and Procházka, M., “Drop coating deposition Raman spectroscopy of dipicolinic acid,” Journal of Raman Spectroscopy 49, 2050–2052 (dec 2018).

[46] Setlow, P. and Christie, G., “What’s new and notable in bacterial spore killing!,” World Journal of Microbiology and Biotechnology 37, 144 (aug 2021).

[47] McCann, K. and Laane, J., “Raman and infrared spectra and theoretical calculations of dipicolinic acid, dinicotinic acid, and their dianions,” Journal of Molecular Structure 890, 346–358 (nov 2008).

[48] Kong, L., Setlow, P., and Li, Y.-q., “Analysis of the Raman spectra of Ca2+-dipicolinic acid alone and in the bacterial spore core in both aqueous and dehydrated environments,” The Analyst 137(16), 3683 (2012).

[49] Malyshev, D., Oberg, R., Dahlberg, T., Wiklund, K., Landström, L., Andersson, P. O., and Andersson, M., “Laser induced degradation of bacterial spores during micro-Raman spectroscopy,” Spectrochimica Acta Part A: Molecular and Biomolecular Spectroscopy 265, 120381 (jan 2022).

[50] Shibata, H., Yamashita, S., Ohe, M., and Tani, I., “Laser Raman spectroscopy of lyophilized bacterial spores,” Microbiology and immunology 30(4), 307–313 (1986).

[51] McDonnell, G. and Russell, a. D., “Antiseptics and disinfectants: activity, action, and resistance.,” Clinical microbiology reviews 12, 147–79 (jan 1999).

[52] Huang, S.-s., Chen, D., Pelczar, P. L., Vepachedu, V. R., Setlow, P., and Li, Y.-q., “Levels of Ca 2+-Dipicolinic Acid in Individual *Bacillus* Spores Determined Using Microfluidic Raman Tweezers,” Journal of Bacteriology 189, 4681–4687 (jul 2007).

[53] Stöckel, S., Schumacher, W., Meisel, S., Elschner, M., Rösch, P., and Popp, J., “Raman Spectroscopy-Compatible Inactivation Method for Pathogenic Endospores,” Applied and Environmental Microbiology 76, 2895–2907 (may 2010).

[54] Zhang, P., Kong, L., Setlow, P., and Li, Y.-q., “Characterization of Wet-Heat Inactivation of Single Spores of Bacillus Species by Dual-Trap Raman Spectroscopy and Elastic Light Scattering,” Applied and Environmental Microbiology 76, 1796–1805 (mar 2010).

[55] Berry, D., Mader, E., Lee, T. K., Woebken, D., Wang, Y., Zhu, D., Palatinszky, M., Schintlmeister, A., Schmid, M. C., Hanson, B. T., Shterzer, N., Mizrahi, I., Rauch, I., Decker, T., Bocklitz, T., Popp, J., Gibson, C. M., Fowler, P. W., Huang, W. E., and Wagner, M., “Tracking heavy water (D2O) incorporation for identifying and sorting active microbial cells,” Proceedings of the National Academy of Sciences of the United States of America 112(2), E194–E203 (2015).

[56] Zhang, M., Hong, W., Abutaleb, N. S., Li, J., Dong, P. T., Zong, C., Wang, P., Seleem, M. N., and Cheng, J. X., “Rapid Determination of Antimicrobial Susceptibility by Stimulated Raman Scattering Imaging of D2O Metabolic Incorporation in a Single Bacterium,” Advanced Science 7(19), 1–14 (2020).

[57] Yang, K., Li, H. Z., Zhu, X., Su, J. Q., Ren, B., Zhu, Y. G., and Cui, L., “Rapid Antibiotic Susceptibility Testing of Pathogenic Bacteria Using Heavy-Water-Labeled Single-Cell Raman Spectroscopy in Clinical Samples,” Analytical Chemistry 91(9), 6296–6303 (2019).

[58] Liu, Y., Ma, Y., Zhang, L., Sun, X., Yang, J., Li, X., Li, F., Chen, R., Zhu, P., Xu, J., and Yang, F., “Singlecell Raman microspectroscopy-based assessment of three intracanal disinfectants’ effect on *Enterococcus faecalis*,” Journal of Raman Spectroscopy 53(5), 902–910 (2022).

[59] Öberg, R., Dahlberg, T., Malyshev, D., and Andersson, M., “Monitoring bacterial spore metabolic activity using heavy water-induced Raman peak evolution,” (Under Review) Analytical Chemistry (2022).

[60] Svoboda, K. and Block, S. M., “Optical trapping of metallic Rayleigh particles.,” Optics letters 19(13), 930–932 (1994).

[61] Ayano, S., Wakamoto, Y., Yamashita, S., and Yasuda, K., “Quantitative measurement of damage caused by 1064-nm wavelength optical trapping of Escherichia coli cells using on-chip single cell cultivation system,” Biochemical and Biophysical Research Communications 350, 678–684 (nov 2006).

[62] Mirsaidov, U., Timp, W., Timp, K., Mir, M., Matsudaira, P., and Timp, G., “Optimal optical trap for bacterial viability,” Physical Review E 78, 021910 (aug 2008).

[63] Zhang, Y., Miao, Z., Huang, X., Wang, X., Liu, J., and Wang, G., “Laser Tweezers Raman Spectroscopy (LTRS) to Detect Effects of Chlorine Dioxide on Individual Nosema bombycis Spores,” Applied Spectroscopy 73(7), 774–780 (2019).

[64] Mohanty, S. K., Rapp, A., Monajembashi, S., Gupta, P. K., and Greulich, K. O., “Comet assay measurements of DNA damage in cells by laser microbeams and trapping beams with wavelengths spanning a range of 308 nm to 1064 nm,” Radiation Research 157(4), 378–385 (2002).

[65] Jockusch, S., Turro, N. J., Thompson, E. K., Gouterman, M., Callis, J. B., and Khalil, G. E., “Singlet molecular oxygen by direct excitation,” Photochem. Photobiol. Sci. 7(2), 235–239 (2008).

[66] Moeller, R., Raguse, M., Reitz, G., Okayasu, R., Li, Z., Klein, S., Setlow, P., and Nicholson, W. L., “Resistance of Bacillus subtilis Spore DNA to Lethal Ionizing Radiation Damage Relies Primarily on Spore Core Components and DNA Repair, with Minor Effects of Oxygen Radical Detoxification,” Applied and Environmental Microbiology 80, 104–109 (jan 2014).

[67] Cybulski, R. J., Sanz, P., Alem, F., Stibitz, S., Bull, R. L., and O’Brien, A. D., “Four Superoxide Dismutases Contribute to Bacillus anthracis Virulence and Provide Spores with Redundant Protection from Oxidative Stress,” Infection and Immunity 77, 274–285 (jan 2009).

[68] Patil, P. V. and Ballou, D. P., “The use of protocatechuate dioxygenase for maintaining anaerobic conditions in biochemical experiments,” Analytical Biochemistry 286(2), 187–192 (2000).

[69] Mesu, J. G., Visser, T., Soulimani, F., and Weckhuysen, B. M., “Infrared and Raman spectroscopic study of pH-induced structural changes of l-histidine in aqueous environment,” Vibrational Spectroscopy 39, 114–125 (sep 2005).

[70] Ashton, L., Johannessen, C., and Goodacre, R., “The Importance of Protonation in the Investigation of Protein Phosphorylation Using Raman Spectroscopy and Raman Optical Activity,” Analytical Chemistry 83, 7978–7983 (oct 2011).

